# KMT2D Loss Promotes Head and Neck Squamous Carcinoma Through Enhancer Reprogramming and Modulation of Immune Microenvironment

**DOI:** 10.1101/2021.09.21.461314

**Authors:** S. Carson Callahan, Margarita Divenko, Praveen Barrodia, Anand K Singh, Emre Arslan, Zhiyi Liu, Jiah Yang, Nazanin Anvar, Moran Amit, Tongxin Xie, Shan Jiang, Jonathan Schulz, Ming Tang, Jeffrey N Myers, Kunal Rai

**Affiliations:** Department of Genomic Medicine, The University of Texas MD Anderson Cancer Center, Houston, TX 77054, USA; Department of Head and Neck Surgery, The University of Texas MD Anderson Cancer Center, Houston, TX 77030, USA; MD Anderson Cancer Center UTHealth Graduate School of Biomedical Sciences, University of Texas MD Anderson Cancer Center, Houston, Texas, USA; FAS informatics Group, Harvard University, Cambridge, MA USA; Department of Molecular and Cellular Biology, Center for Brain Science, Harvard University and Howard Hughes Medical Institute, Cambridge, MA USA

## Abstract

Head and neck squamous cell carcinoma (HNSCC) is the sixth most common cancer worldwide, with 5-year survival of ∼50%. Genomic profiling studies have identified important somatic mutations in this disease which presents an opportunity for precision medicine. We demonstrate that KMT2D, a histone methyltransferase harbors somatic mutations in ∼17% of HNSCC and is associated with 2-year recurrence in TCGA data. Consistent with algorithmic prediction of bring a driver tumor-suppressor event, its loss results in larger oral tumors in immune-proficient orthotopic models. Mechanistically, we find that KMT2D knockdown or KMT2D mutation causes loss of H3K4me1-marked enhancers harboring IRF7/9 binding sites, which is known to regulate interferon signaling. Indeed, KMT2D loss in human and murine cell lines deregulated transcriptional levels of cytokine expression and impacted numerous immune signaling pathways, including interferon signaling. Consistently, Kmt2d knockdown in murine tumors exhibited decrease in IFN*γ*-producing effector T cells and an increase in T-cells with an exhausted phenotype. Epistasis experiments showed that exogenous treatment with IFN*γ* abrogated the increased tumor growth in Kmt2d-deficient oral tumors. Together, these results support the role of KMT2D as a tumor suppressor in HNSCC that regulates the tumor microenvironment by modulating H3K4me1-marked enhancers controlling interferon signaling.

## INTRODUCTION

Head and neck squamous cell carcinoma (HNSCC) is the sixth most common cancer worldwide, with an estimated 600,000 new cases and 355,000 deaths attributable to the disease each year (Argiris et al., 2008; Zhou et al., 2016). The 5-year survival for HNSCC is approximately 50% and has remained relatively unchanged for decades, owing largely to factors such as late stage at initial presentation and high rates of primary tumor recurrence (Bonner et al., 2006; Pickering et al., 2013). Most efforts in understanding HNSCC and developing targeted therapeutics have focused on genomic characterization, such as exome sequencing and copy number aberrations. This has resulted in the identification of common alterations, such as *TP53* mutations and alterations in the *p16* locus at 9p21(Cancer Genome Atlas, 2015; Forastiere et al., 2001; Zhou et al., 2016). To date, there are notably few targeted therapies in HNSCCC, indicating additional work is needed to identify biomarkers of response, novel therapeutic compounds, and mechanisms of HNSCCC tumorigenesis and progression(Bonner et al., 2006; Gougis et al., 2019). Since the landmark paper in 1996 by Leach et al. (Leach et al., 1996), cancer immunotherapy has also risen in prominence as an approach to treating multiple types of tumors. In the case of HNSCC, studies demonstrate that checkpoint blockade in HNSCC can provide clinical benefit; however, only 20-30% of patients see survival benefit with PD-1 or PDL-1 checkpoint blockade, indicating needs for patient stratification for existing therapies and discovery of novel ways to modulate the immune system for therapeutic benefit (Burtness et al., 2019; Ferris et al., 2016).

Importantly, the tumor environment is not composed solely of tumor cells, but a wide array of different stromal cells, ranging from fibroblasts to immune cells. In all cases, the behavior of the tumor microenvironment (TME) and the cancer cells are intimately linked by crosstalk between the two compartments in the form of cytokines, growth factors, metabolites, and other signaling molecules (Marks et al., 2016; Pitt et al., 2016). In cancer, as in infection, the first line of immune response to altered homeostasis lies in the innate immune system (granulocytes, macrophages, dendritic cells, etc.). In addition to the innate immune system’s ability to directly contribute to killing tumorigenic cells (e.g. via NK cell mediated death and remodeling of the TME matrix via secreted matrix metalloproteinases), one of its chief functions is transducing the response signal to the adaptive arm of the immune system via specialized antigen presenting cells (APCs) (Janeway and Medzhitov, 2002; Medzhitov and Janeway, 2002; Pitt et al., 2016). In particular, upregulation of costimulatory signaling molecules such as CD80 and CD86 by the APCs primes CD8+ and CD4+ T cells for clonal expansion upon recognition of foreign antigen, allowing the adaptive immune system to ultimately clear the immediate threat to the host (Pitt et al., 2016). This transition to an adaptive response is considered to be critical in limiting tumor growth and progression, as evidenced by correlations between the presence of these cells and positive clinical outcomes (Fridman et al., 2012; Pitt et al., 2016). However, recent research has demonstrated the TME can also have immunosuppressive features depending upon the presence of known subsets of cells, like myeloid-derived suppressor cells (a mixture of immature myeloid cells correlated with cancer progression) and T-regulatory cells (a subset of CD4+ cells which inhibit tumoricidal activity of CD8+ T cells and NK cells), or differentiation states of other cells, like macrophages (where the M1 state is associated with tumoricidal activity, and the M2 state is associated with immunosuppressive activity) (Joyce and Pollard, 2009; Klymenko and Nephew, 2018; Pitt et al., 2016). The ability of tumor cells to alter their cytokine expression, metabolism, and antigen presentation machinery provides a key link between the genetic and epigenetic status of the tumor cells and the resulting TME, indicating a need to better understand this interaction and how the tumor cell state affects recruitment of pro- or anti-tumorigenic immune subsets.

In recent years, research has demonstrated that epigenetic changes in tumors can have profound impacts on tumor characteristics such as growth rates and invasive potential as well as therapeutic response (Baylin and Jones, 2016; Castilho et al., 2017). This has, in part, been attributed to epigenetic remodeling in subsets of the tumor cells themselves, leading to generation of a cancer stem cell (CSC) population that contributes to tumor evolution, heterogeneity, and drug resistance (Easwaran et al., 2014; Gupta et al., 2011; Toh et al., 2017). In HNSCC, there is evidence that loss of H3K9ac may be a marker for cisplatin resistance, as well as many papers demonstrating overexpression of histone deacetylase (HDAC) proteins in advanced stage tumors and leading to oncogenic properties such as invasion and increased proliferation rates (Almeida et al., 2014; Castilho et al., 2017). The effect of epigenetic alterations in tumors is still an area of active research, with many studies focusing on directly targeting proteins responsible for placing and removing histone modifications, DNA methylation, and other epigenetic remodeling processes (Baylin and Jones, 2016; Toh et al., 2017). In the case of HNSCC, most success has been seen with HDAC inhibitors and inhibitors of DNA methyltransferases, particularly when used in combination with other chemotherapeutic agents (Castilho et al., 2017).

In this study, we demonstrate that KMT2D, a histone lysine methyltransferase mutated in approximately 18% of TCGA HNSCC tumors (Cancer Genome Atlas, 2015), functions as a tumor suppressor in HNSCC. Further, we demonstrate that loss of KMT2D function affects tumor intrinsic immune response pathways by disrupting the enhancers of relevant genes, ultimately altering the tumor-TME crosstalk, resulting in an “immune cold” TME and larger tumor growth *in vivo*.

## RESULTS

### KMT2D Functions as a Tumor Suppressor In HNSCC

To discover key epigenetic drivers of HNSCC, we first explored the TCGA HNSCC dataset. Of note, we restricted our analysis to HPV (-) tumors (hereafter, simply “HNSCC”), as HPV itself can cause changes in gene expression and chromatin structure and confound interpretation of the data (Kelley et al., 2017). After removal of samples with detectable expression of HPV transcripts or HPV integration, 444 HPV (-) samples were carried forward for further analysis. Cancer “driver” mutations are a class of mutations that confer selective advantage to a cancer cell and its progeny and are heavily implicated in tumorigenesis. This contrasts with “passenger” mutations, which confer no selective advantage and simply remain in the genome of the cancer cell (Martinez-Jimenez et al., 2020; Stratton et al., 2009). To evaluate this hypothesis, we used the OncodriveFML framework to evaluate all the coding regions in the TCGA HNSCC whole exome sequencing (WES) dataset for their “driving” potential (Mularoni et al., 2016). KMT2D was highly ranked amongst all predicted driver genes, which included other known HNSCC drivers, such as TP53, CDKN2A, NOTCH1, and PIK3CA (Cancer Genome Atlas, 2015; India Project Team of the International Cancer Genome, 2013; Pickering et al., 2013; Stransky et al., 2011) **(Figure 1A)**. Interestingly, it was also the most highly ranked epigenetic modifier (histone acetyltransferases and deacetylases, histone methyltransferases and demethylases, and DNA modifiers) in the dataset. We found this particularly intriguing, as recent work has demonstrated KMT2D is an early mutagenic event in human bladder and human HNSCC, furthering the notion that KMT2D mutations are early driver events in some human cancers (Lawson et al., 2020; Li et al., 2020; Veeramachaneni et al., 2019).

**Figure 1:**
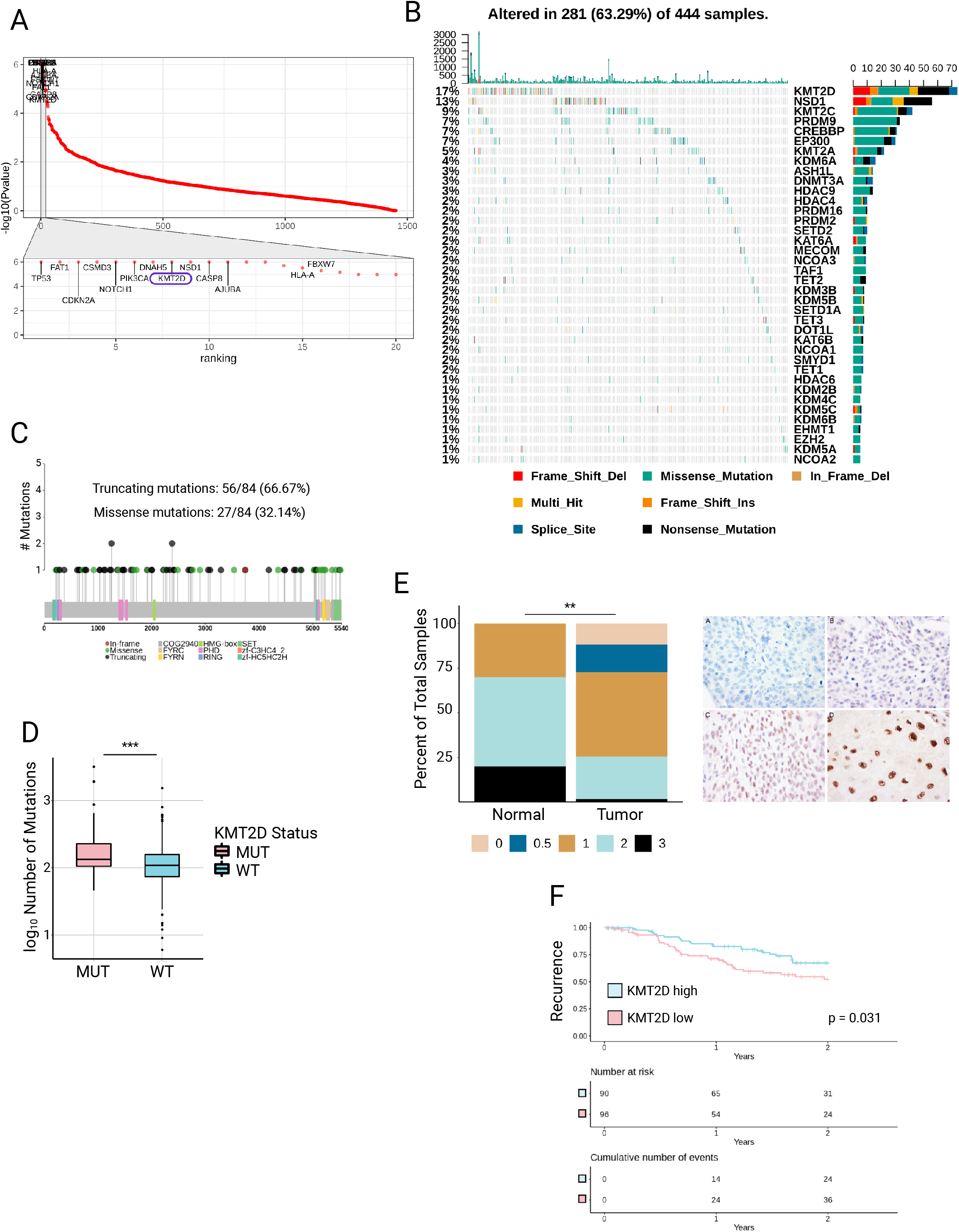
KMT2D Functions as a Tumor Suppressor in HNSCC. **a**. Ranking of “driving” mutations across all CDS regions in the HPV (-) HNSCC TCGA WES data. The bottom box represents a zoomed in view of the top hits with KMT2D circled in purple. **b**. Oncoplot of epigenetic modifiers (histone acetyltransferases and deacetylases, histone methyltransferases and demethylases, and DNA modifiers) in TCGA HNSCC. **c**. Lollipop plot of all KMT2D mutations from **(b)**. Green dots are missense mutations, black dots are truncations, and red dots are in-frame mutations. The SET domain is represented by the green box at the C terminal end of the protein. **d**. Boxplot of total number of mutations in KMT2D mutant and KMT2D WT samples in TCGA HNSCC (***, *p* < 0.001, Mann-Whitney *U* test). **e**. Barplot demonstrating staining intensity scores from KMT2D IHC performed on a human HNSCC TMA slide (left)(**, *p* < 0.01, Mann-Whitney *U* test) and representative examples of staining intensity scores (right). **f**. Two-year recurrence free survival of KMT2D low expressing (pink) and KMT2D high expressing (light blue) patients in TCGA HNSCC (*p =* 0.031, log-rank test).

To further explore the epigenetic mutational landscape in HNSCC, we examined the mutation rates of known epigenetic modifiers and found KMT2D to be mutated in 17% of HNSCC, making it the most highly mutated epigenetic modifier and one of the most highly mutated genes in HNSCC overall **(Figure 1B, S1A)**. Previous biochemical investigations of KMT2D function indicate that not all mutations have functional consequences for H3K4 methylation, with disruptions in methyltransferase activity only occurring if mutations are truncating or disrupt the C-terminal SET domain (Zhang et al., 2015). Using the TCGA WES dataset, we investigated the different classes of mutations and their distribution across the gene. Interestingly, we found that most mutations are classed as truncations (56/84, 66.67%), and two missense mutation clusters occur in the C-terminal end of the protein at the methyltransferase (SET) domain and other functional domains **(Figure 1C)**, suggesting these mutations have functional consequences and contribute to HNSCC development. Additionally, we found that samples with KMT2D mutations have a significantly higher overall mutation burden than KMT2D WT samples (*p* < 0.001, **Figure 1D**), further supporting KMT2D loss as a key event in HNSCC development.

To confirm our findings at the protein level, we performed IHC against KMT2D on a human HNSCC TMA containing both tumor and normal head and neck tissue. The panel was scored by a medical pathologist using a 0 to 3 system **(Figure 1E)**, with higher numbers indicating more intense signal. We compared the intensity of signal in samples derived from primary tumor and normal tissue and found that the distribution of staining scores differed significantly (*p* < 0.01), with tumor samples displaying lower staining scores than normal tissue **(Figure 1E)**, consistent with a characterization of KMT2D as a tumor suppressor in HNSCC.

We next assessed the relationship between KMT2D and clinical outcomes using the TCGA HNSCC clinical dataset. We divided samples into quartile groups based on KMT2D expression in the TCGA RNA-seq dataset and compared the top and bottom quartile groups. Using these data, we discovered that patients in the bottom quartile of KMT2D expression had significantly worse 2-year recurrence rates than patients in the top quartile of KMT2D expression (*p* < 0.05, **Figure 1F**). This finding is particularly interesting in the case of HNSCC, as approximately 50% of HNSCC recurrences are within the first 2 years of remission, and median survival of patients with recurrence is less than 1 year (Argiris et al., 2008; Vermorken et al., 2008). Together, these findings support the role of KMT2D as a tumor suppressor in HNSCC that contributes to clinically relevant patient outcomes.

### KMT2D Regulates Response to Interferon and Cytokine Signaling

KMT2D is a histone lysine methyltransferase with a variety of critical functions, ranging from regulation of metabolism in cancer to control of skin homeostasis in conjunction with p63 (Koutsioumpa et al., 2019; Lin-Shiao et al., 2018; Maitituoheti et al., 2020). To study the effects of KMT2D loss in HNSCC, we first generated KMT2D knockdown (KD) in the established HNSCC cell line HN30 and performed RNA-seq. Differential expression analysis of KMT2D KD vs. non-targeted control (NT) displayed widespread disruption of the HNSCC transcriptome with many upregulated and downregulated genes. Of note, we observed significant downregulation of many genes involved in mediating interferon response, such as the IRF family of transcription factors, and several downstream target genes, such as the IFIT family of proteins, suggesting KMT2D may play a role in modulating tumor cell responses to immune signaling **(Figure 2A)**. Indeed, pathway analysis of our KMT2D KD RNA-seq data showed significant negative enrichment for numerous immune-related pathways, particularly those involved in interferon response and JAK-STAT signaling **(Figure 2B)**. We next investigated if similar patterns are observed in the KMT2D-mutant human tumors by performing differential expression analysis on the TCGA RNA-seq dataset. Using the quartile groups previously generated, we compared KMT2D low expressing samples to KMT2D high expressing samples. Similar to our results in the KMT2D KD system, we observed disruption of numerous members of the JAK-STAT pathway and IRF family of transcription factors **(Figure 2C)**. Pathway analysis of this data demonstrated significant negative enrichment in immune-related pathways, in addition to a significant positive enrichment in metabolic pathways, particularly those related to oxidative energy-producing processes, such as oxidative phosphorylation and fatty acid metabolism **(Figure 2D)**. These findings are consistent with previous findings in melanoma and lung tumors, indicating KMT2D loss may play similarly important roles in multiple tumor types (Alam et al., 2020; Maitituoheti et al., 2020).

**Figure 2:**
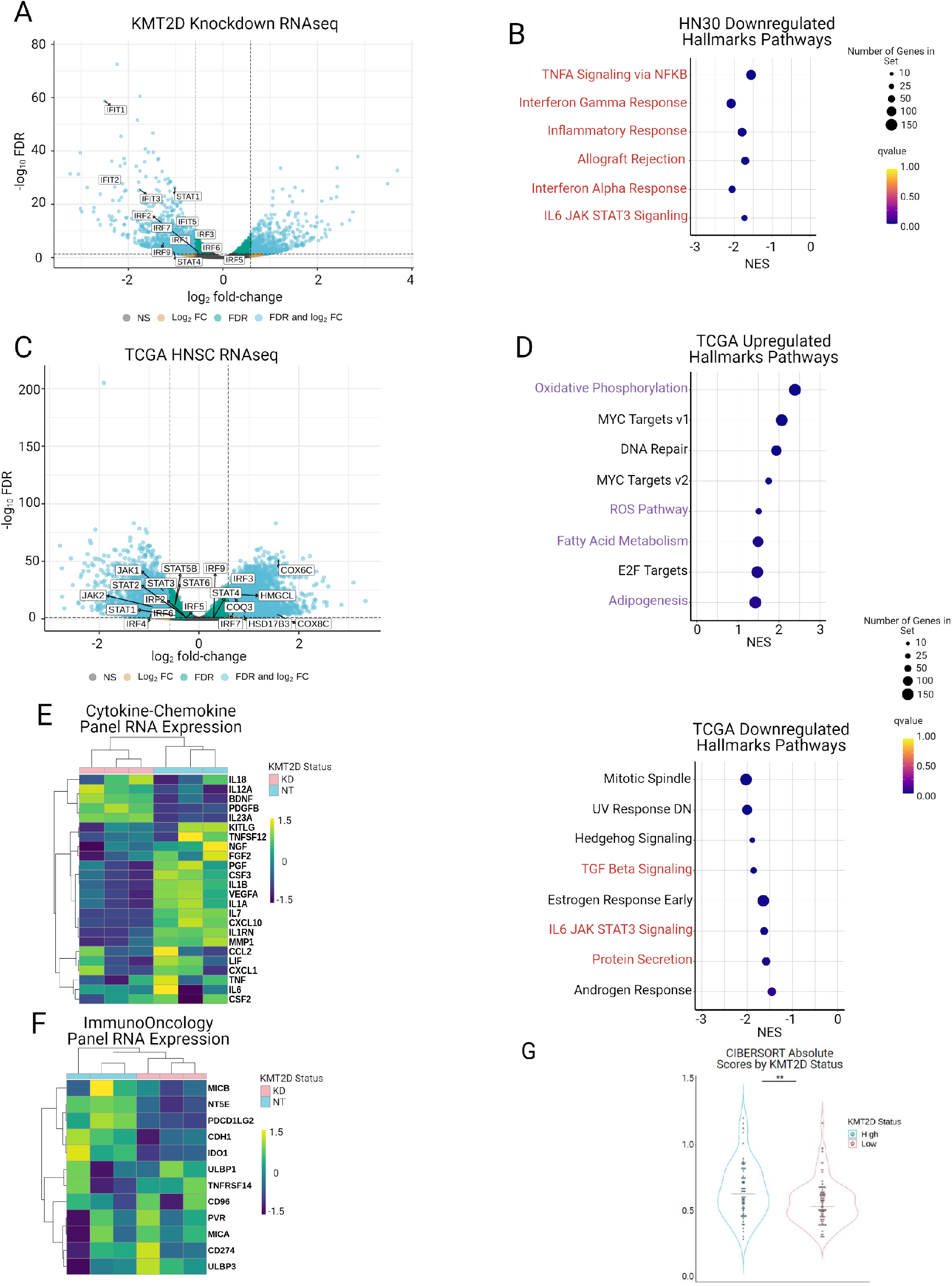
KMT2D Regulates the Expression of Tumor-Intrinsic Immune Response Genes. **a**. Volcano plot demonstrating differentially expressed genes in the HN30 cell line after knockdown with shKMT2D (FDR < 0.05, abs(fold-change) > 1.5). **b**. Dotplot visualizing top downregulated Hallmarks Pathways in HN30 shKMT2D per GSEA. Pathways highlighted in red are associated with immune signaling. **c**. Volcano plot demonstrating differentially expressed genes in KMT2D low TCGA HNSCC samples compared to KMT2D high TCGA HNSCC samples (FDR < 0.05, abs(fold-change) > 1.5). **d**. Dotplots visualizing top upregulated (top) and downregulated (bottom) Hallmarks Pathways in KMT2D low expressing samples from TCGA HNSCC. Pathways highlighted in purple are associated with metabolic and oxidative processes; pathways highlighted in red are associated with immune signaling. Hierarchical clustering of cytokine-chemokine **(e)** and immuno-oncology **(f)** gene panels based on HN30 shKMT2D RNA-seq data performed using Ward’s method. **g**. Violin plots demonstrating CIBERSORT absolute scores in KMT2D low expressing and KMT2D high expressing groups from the TCGA HNSCC data (**, *p* < 0.01, Mann-Whitney *U* test).

Since tumor cells secrete numerous factors (cytokines, growth factors, etc.) that allow for crosstalk between tumor cells and the TME and JAK-STAT signaling is known to regulate the production of similar factors, we reasoned KMT2D loss might alter the profile of factors secreted by tumor cells (Klymenko and Nephew, 2018; Seif et al., 2017). To investigate this hypothesis, we performed hierarchical clustering on the KMT2D KD RNA-seq data, selecting genes from lists from pre-established cytokine/chemokine and immuno-oncology panels that were also expressed in our tumor cell line. Clustering results demonstrated substantial alterations in the expression patterns of both the cytokine-chemokine panel **(Figure 2E)** and the immuno-oncology panel **(Figure 2F)**. Of note, the cytokine-chemokine panel demonstrated significant decreases in the expression of CXCL10, a chemokine responsible for chemoattraction of numerous immune cell types, CSF3, a cytokine that stimulates proliferation and survival of macrophages and neutrophils, and IL7, a cytokine important for the development and survival of B and T cells **(Figure 2E)** (ElKassar and Gress, 2010; Liu et al., 2011). In the immuno-oncology panel, we noted significant downregulation of PDCD1LG2, also known as PDL2, but not CD274, also known as PDL1 **(Figure 2F)**. To investigate if KMT2D loss results in an altered immune TME, we utilized the CIBERSORT algorithm to deconvolve the TCGA RNA-seq data into its respective immune populations (Newman et al., 2015). We compared the total absolute immune scores of KMT2D low to KMT2D high samples and found that, indeed, KMT2D low samples had a significantly lower total absolute immune score (*p* < 0.01) **(Figure 2G)**. Together, these findings suggest KMT2D loss results in a significant downregulation of numerous genes involved in the tumor cell intrinsic response to interferon and cytokine signaling, leading to an alteration in the cytokine/chemokine and cell surface marker expression profiles of the tumor cells themselves.

### An Orthotopic Model of KMT2D Loss Demonstrates Larger Tumor Volume and an Altered Immune Repertoire

To study the effect of KMT2D *in vivo*, we utilized the MOC1 syngeneic mouse model of oral cavity tumors, which has previously been used in immune TME interaction studies and in other genomic studies (Judd et al., 2012a; Judd et al., 2012b; Onken et al., 2014). Using the established mouse oral cavity tumor MOC1 cell line, we generated doxycycline inducible Kmt2d knockdown (shKmt2d) and control (shNT) cells **(Figure 3A)**. These cells were implanted into the oral tongue of C57BL/6 mice, which were split into 4 categories: shKmt2d Dox (-), shKmt2d Dox (+), shNT Dox (-), and shNT Dox (+). Mice were monitored for 4 weeks, at which point samples were collected for further processing and analysis. Analysis of endpoint tumor volumes revealed significantly larger tumor volumes in shKmt2d Dox (+) mice compared to shNT Dox (+) mice (*p* < 0.05), supporting the role of KMT2D as a tumor suppressor in HNSCC **(Figure 3B)**.

**Figure 3.**
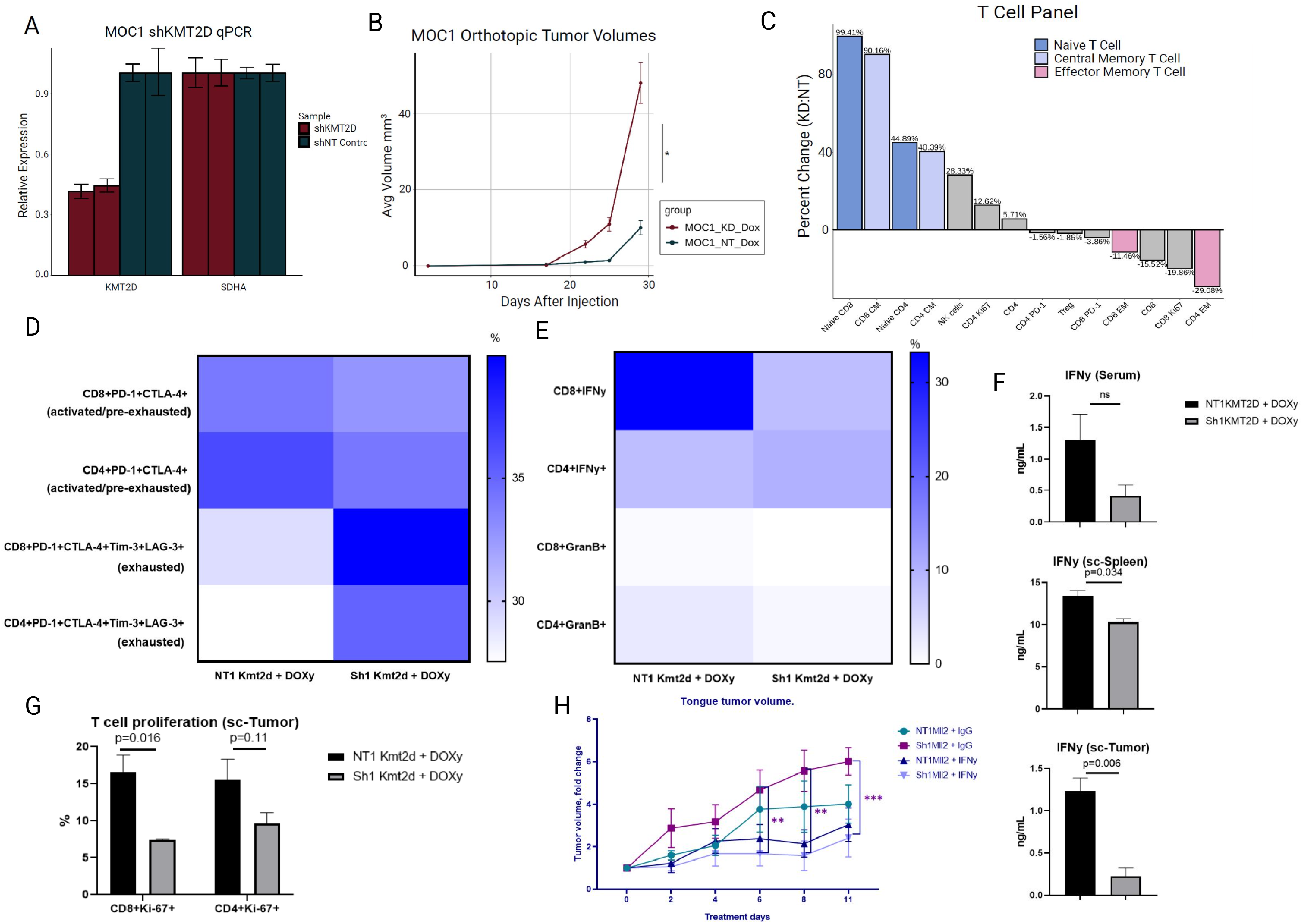
An Orthotopic Model of KMT2D Loss Demonstrates Larger Tumor Volume and an Altered Immune Repertoire. 4-6 weeks old female mice on C57BL/6 background were orthotopically injected with either ShNT or ShKmt2d MOC1 cell lines. **a**. q-PCR data illustrating substantial decrease of mRNA level of Kmt2d in ShKmt2d MOC1 cell line. **b**. Graph demonstrating kinetics of tongue tumor volume change after orthotopic injection (*n=5*). Flow cytometry analysis to characterize T cells **c**. The difference in the main T cells populations shown as percent change between two experimental groups. Single cell-suspension isolated from tongue tumor was stimulated for 4 h in the presence of phorbol 12-myristate-13-acetate, ionomycin, and brefeldin A and subjected to intracellular and nuclear flow cytometry staining: Heat maps demonstrating the percentage of **d**. surface co-stimulatory markers and **e**. IFNy and Granzyme B cytokines on total CD4+ and CD8+ T cells. **g**. Percentage of CD4+Ki-67+and CD8+Ki-67+ T cells out of total CD4+ and CD8+ T cells (*n=5*). **f**. The level of IFNy detected by ELISA in serum and supernatant of cultured overnight in the presence of 2.5 ug agonistic anti-CD3 and anti-CD28 antibodies and 1 ug/mL of LPS single cells isolated from tongue tumor and spleen (*n=5*). **h**. Graph illustrating tongue tumor fold change after orthotopic injection of either ShNT or ShKmt2d MOC1 cell lines and IFNy i.p. administration (1 ug per mouse) into 4-6 weeks old female mice on C57BL/6 background (*n=4*). p-value in **b, f, g, h** was calculated using two-sided unpaired Student’s *t*-test; *n* represents the number of experimental animals per group. Data are representative of one individual experiment and are presented as mean ± SEM and mean ± SD.

We next investigated the immune TME of shNT Dox (+) and shKmt2d Dox (+) tumors by FACS analysis which substantial differences in adaptive immune response. Particularly, we detected a 44.8% increase in CD4+ naive T cells as well as a shift in CD4+ memory cell distribution, with a 40% increase in CD4+ central memory cells and a 29% decrease in a CD4+ effector memory cells **(Figure 3C)**. Interesting, these same trends were observed in the CD8+ T cell populations, with a 99% increase in CD8+ naive T cells, a 90% increase in CD8+ central memory cells, and an 11% decrease in CD8+ effector memory cells **(Figure 3C)**.

To further elaborate on immune response alteration in TME upon Kmt2d loss in tumor cells, we performed additional FACS analysis of tongue tumor specimens in doxycycline inducible MOC1 *in vivo* model described above. The shift in T cells’ populations toward more naive stage and decrease of effector phenotype in both CD4+ and CD8+ T cells **(Figure 3C)** made us investigate the proliferation, activation, and cytotoxic features of T cells in depth. Both CD4+ and CD8+ T cells in mice bearing Kmt2d KD tumors have decreased level of proliferation marker Ki-67 (**Figure 3G)**. Additionally, to investigate the activation status of T cells we detected the surface expression level of co-stimulatory molecules such as PD-1, CTLA-4, Tim-3, LAG-3. FACS analysis revealed that double positive PD-1+ CTLA-4+ T cells populations, mostly comprised of fully activated and or reversible pre-exhausted T cells, were decreased in Kmt2d KD group **(Figure 3D)**, whereas PD-1+ CTLA-4+ Tim-3+ LAG-3+ positive irreversibly exhausted T cells populations were increased in the Kmt2d KD group **(Figure 3D)**. Besides, based on the intracellular flow cytometry staining, CD8+ T cells lost their capacity to produce IFN*γ* in TME of Kmt2d KD tumors **(Figure 3E)**. These data suggest that T cells in the setting of Kmt2d loss are characterized by less proliferative and activated phenotype, lose their cytotoxic activity, demonstrated by IFN*γ* level decrease, and have overexpression of exhaustion markers.

The role of IFN*γ* in anti-tumor immunity has been well established (Ayers et al., 2017; Briesemeister et al., 2011; Tau et al., 2001). To test the global and local levels of IFN*γ*, we performed ELISA analysis of serum and activated cells isolated from spleen and tongue tumors. Expectedly, we detected significant decrease of IFN*γ* at the local tissue site as well as at the secondary lymphoid organ in the settings of Kmt2d loss **(Figure 3F)**. The global level of circulating IFN*γ* tested on serum samples did not reveal any differences between two experimental groups **(Figure 3F)**. The substantial decrease of IFN*γ* can be due to loss of proliferative cytoxic T cells populations upon Kmt2d deficiency. Moreover, the lack of IFN*γ* can further leads to reduced T cells’ differential potential due to disrupted IFN*γ* autocrine loop in T cells. To assess if supplementation of IFN*γ* could rescue the observed reduction of anti-tumor immunity and could help to decrease the tumor burden, we conducted *in vivo* studies where we utilized the same doxycycline inducible MOC1 system and orthotopically implanted shNT and shKmt2d tumors in C57BL/6 mice. At the 12^th^ day after tumor establishment mice were given IFN*γ* via intraperitoneal injections every other day. By the 6^th^ day of treatment mice bearing, Kmt2d KD tumors started benefiting from IFN*γ* supplementation as indicated by reduced tumor volumes **(Figure 3H)**. These results additionally demonstrate that Kmt2d loss interferes with immune cells signaling which leads to downregulation of anti-tumor capacity by immune cells, primarily T cells, and affects the local level of IFN*γ* which further exacerbates the observed phenotype. Together, these findings provide evidence that Kmt2d loss in the tumor leads to reduction of proliferated IFN*γ*-producing effector T cells, which ultimately leads to decrease of IFN*γ* level at the TME which in turn increases tumor burden.

### Enhancer reprogramming in KMT2D-deficient head and neck cancer cells

To further gain mechanistic insights into how KMT2D may modulate the immune microenvironment and because of its histone methyltransferase avtivity, we investigated its role in modulating the epigenome in HNSCC. To define basal epigenetic states, we performed chromatin immunoprecipitation followed by sequencing (ChIP-seq) on a panel of 6 histone marks (H3K4me1, H3K4me3, H3K27ac, H3K27me3, H3K9me3 and H3K79me2) in HN30 shNT control and HN30 shKMT2D knockdown cell lines. Using ChromHMM, we defined “chromatin states”, i.e. different combinations of histone marks, and their genome-wide binding patterns. These states are then annotated based on their emission profiles generated by the ChromHMM model to obtain a complete picture of the entire genome’s activity **(Figure 4A)**. Our analysis reveals that, after KMT2D knockdown, there is a loss of signal in states containing the H3K4me1 mark (represented by a transition to the E11 “Low” state in HN30 shKMT2D) **(Figure 4B)**. Given the literature support for the role of KMT2D as a modulator of H3K4me1, we focused our analysis on alterations in chromatin states containing this mark (Froimchuk et al., 2017; Lee et al., 2013; Lin-Shiao et al., 2018; Local et al., 2018; Maitituoheti et al., 2020; Rao and Dou, 2015; Wang et al., 2016). Based on the annotation of these states, we were particularly interested in the loss of signal in states E09 and E10, as these represent enhancer states characterized by either high H3K4me1 signal (E09) or high H3K27ac signal (E10) **(Figure 4A-B)**. Comparing H3K4me1-marked regions in HN30 shNT to HN30 shKMT2D, we discovered that shNT had 4866 unique H3K4me1 peaks and shKMT2D had 2169 unique peaks **(Figure 4C)**. This demonstrates that loss of KMT2D results in an overall loss of H3K4me1 signal, consistent with its function as a writer of the H3K4me1 histone mark. To correlate this loss of H3K4me1 signal with changes in gene expression, we linked downregulated genes to their putative enhancers using the Cao et al. data, which leverages a model built from 935 samples from a wide variety of primary human cells, tissues, and cell lines to generate enhancer-target networks (Cao et al., 2017). We then generated H3K4me1 signal enrichment plots across these putative enhancers, which demonstrated overall signal loss after KMT2D knockdown **(Figure 4D)**. Curious as to which factors might be affected by this loss of enhancer signal, we performed motif enrichment analysis on those peaks unique to the shNT condition (i.e. enhancers lost in the shKMT2D condition), which revealed an enrichment for IRF factor binding **(Figure 4D)**. IRF factors are involved in regulating immune signaling (Jefferies, 2019; Tamura et al., 2008; Yanai et al., 2012), which is consistent with our previous analysis of RNA-seq data **(Figure 2A-F)**, supporting the notion that KMT2D loss alters the immune response in tumor cells by affecting enhancer activity. To expand on this analysis, we performed H3K4me1 ChIP-seq on a panel of unmodified HNSCC cell lines. Our dataset consists of 5 cell lines with functional KMT2D mutations (stopgain, frameshift deletion, or missense mutations in C-terminal domain regions) and 5 cell lines with WT KMT2D (**Supplementary Table 1**). To reduce possible background effects of other co-existing mutations, KMT2D WT cell lines were selected based on the similarity of their binarized WES mutation profiles using hierarchical clustering based on Jaccard distance. We next performed a differential binding analysis to investigate the effect a natural KMT2D mutation had on H3K4me1, and this analysis revealed that all regions identified as differentially bound (*p* < 0.005 and abs(fold-change) > 1.5) were downregulated in the KMT2D mutant condition compared to KMT2D wildtype **(Figure 4F)**. To investigate the function of these lost enhancer regions, we again leveraged the Cao et al. data to link these regions to their putative genes (Cao et al., 2017). Pathway analysis for these genes demonstrated an enrichment for immune signaling activity, indicating KMT2D mutation mediated enhancer alterations deregulate immune genes in HNSCC **(Figure 4G)**. Overall, epigenomic data supports the role of KMT2D in regulating H3K4me1-marked enhancers that function as regulators of genes involved in immune signaling pathways.

**Figure 4:**
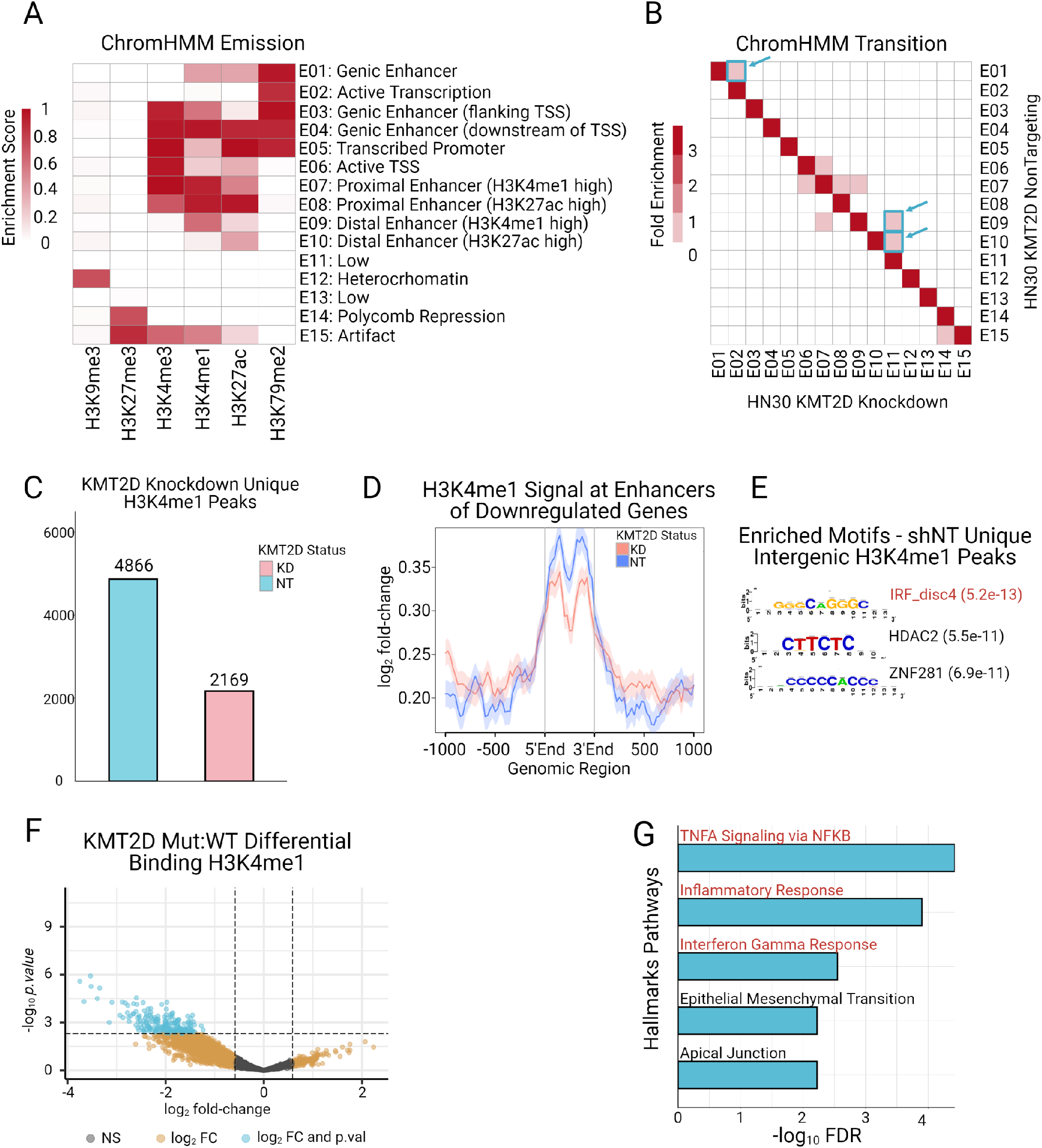
KMT2D Modulates H3K4me1-marked Enhancers for Immune Response Genes in HNSCC. **a**. Annotated 15-state ChromHMM model in HN30 shKMT2D and HN30 shNT. Functional annotations are listed along the Y-axis; histone marks used to generate the model are listed along the X-axis. Signal from any particular histone mark in any given state is indicated by red shading. **b**. Chromatin state transitions from HN30 shNT (Y-axis) to HN30 shKMT2D (X-axis). Boxes highlighted in blue and marked with arrows indicate transitions that involve loss of H3K4me1 after KMT2D knockdown. **c**. Barplot visualizatig the number of unique peaks in both the HN30 shNT (light blue) and HN30 shKMT2D (pink) conditions. **d**. H3K4me1 enrichment across all putative enhancers of genes downregulated after KMT2D knockdown. **e**. ENCODE database motif enrichment in intergenic H3K4me1 peaks unique to HN30 shNT (e-value reported in parentheses next to each motif); motifs highlighted in red are associated with immune factors. **f**. Differentially bound H3K4me1-marked regions in a panel of KMT2D mutant and KMT2D wildtype HNSCC cell lines (*p* < 0.005, abs(fold-change) > 1.5). **g**. Hallmarks Pathways enriched in genes linked to regions significantly downregulated in **(f)**; pathways highlighted in red are associated with immune signaling.

## DISCUSSION

Overall, our results suggest that KMT2D promotes head and neck cancer via reprogramming of immune microenvironment by deregulation of enhancers for immune genes. We present a model wherein, in normal epithelial cells, KMT2D cooperates with IRF7/9 to activate enhancers for interferon response genes including important cytokines which remodel the tumor microenvironment to eventually activate T cells for tumor suppression.

KMT2D is highly mutated across many different cancer types (Kantidakis et al., 2016) (Dhar and Lee, 2021; Kandoth et al., 2013; Rao and Dou, 2015). Previous whole exome sequencing studies of head and neck tumors revealed nonsense and missense mutations in KMT2D in ∼15% of cases (Cancer Genome Atlas, 2015). Our data suggests that this is the most significantly mutated epigenetic protein in this disease and is likely to be a driver event. This provides an opportunity to develop biomarker-driven therapeutic approach for this set of patients. Our previous work revealed dependency of KMT2D-deficient lung cancers and melanoma on glycolysis inhibition and on IGF1 signaling (Alam et al., 2020; Maitituoheti et al., 2020). This study reveals reprogramming of TME as a novel aspect of KMT2D-deficient HNSCCC. Future studies will be needed to examine relationships between metabolic abnormalities and immune reprogramming observed in the KMT2D-deficient tumors. Furthermore, our data also suggests that KMT2D-deficient cancers may be responsive to an immunomodulatory agent, such as IFN*γ*. Future studies could examine whether a combination of immunotherapy and metabolic therapy may be efficacious to reduce growth of KMT2D-mutant tumors.

Our data also demonstrates enhancer reprogramming in KMT2D-deficient head and neck cancers. This is consistent with our previous studies where similar reprogramming of H3K4me1-marked enhancer states were observed in KMT2D-deficient cells (Maitituoheti et al., 2020). Together, these findings solidify KMT2D as the major guardian of enhancer states in cancer cells. This necessitates novel approaches to target loss of enhancer function in this subset of tumors. Specific epigenetic modulators, such as HDAC, could be effective (Alam et al., 2020), however, systematic testing in relevant in vivo models is needed to test this. In summary, our study identifies modulation of tumor microenvironment as an important mechanism in KMT2D-mutant head and neck cancers.

## MATERIALS AND METHODS

### Cell culture

The naturally occurring human HNSCC cell lines HN30, MDA686TU, MDA686LN, MDA886LN, MDA1586, PCI24, UMSCC11A, UMSCC22B, JHU029, and HN5 were acquired and characterized as previously described (Zhao et al., 2011). All cell lines except JHU029 and HN5 were cultured in DMEM supplemented with 10% FBS, L-glutamine, sodium pyruvate, nonessential amino acids, vitamins, and 1% penicillin-streptomycin. Cell line JHU029 was cultured in RPMI 1640 supplemented with 10% FBS, L-glutamine, sodium pyruvate, nonessential amino acids, and 1% penicillin-streptomycin. Cell line HN5 was cultured in DMEM/F12 supplemented with 10% FBS, L-glutamine, and 1% penicillin-streptomycin. The MOC1 cell line was provided by R. Uppaluri (Washington University School of Medicine, St. Louis, Missouri, USA) (Judd et al., 2012b) and cultured in a 2:1 mixture of IMDM and Ham’s F-12 nutrient mixture supplemented with 5% Tet-free FBS, 1% penicillin-streptomycin, 2.5mg insulin, 20µg hydrocortisone, and 2.5µg EGF. HEK293T cells (ATCC^®^ CRL-3216™) were cultured in DMEM supplemented with 10% FBS and 1% penicillin-streptomycin. All cell lines were cultured at 37 °C in an atmosphere of 5% CO_2_.

Human KMT2D knockdown experiments were performed in the HN30 cell line using KMT2D-specific shRNAs described previously (Maitituoheti et al., 2020). Knockdown experiments for the mouse MOC1 cell line were done using the SMARTvector Inducible Lentiviral shRNA system from Horizon Discovery: V3SM11253-233092640 (sh-KMT2D) and VSC11658 (sh-nontargeting). All shRNA vectors were packaged into lentivirus using HEK293T cells and transduced into target cells using 8ug/ml polybrene. After transduction, cells were selected using 1.5 ug/ml puromycin for 72 hours.

### TCGA Analysis

For our TCGA analyses, WES, RNA-seq, and clinical data were all downloaded from Firebrowse (http://firebrowse.org/). To determine if samples were HPV (+), we used data from a recent comprehensive analysis of HNSCC TCGA data (Nulton et al., 2017). Samples with evidence of HPV infection from any strain were excluded from further analysis.

Analysis of mutations in the TCGA WES data were carried out using the Maftools R package (Mayakonda et al., 2018). Lollipop plots were generated using notations taken from cbioportal (https://www.cbioportal.org/). The same HPV (+) sample removed MAF file was used as input for OncoDriveFML mutation analysis using the package’s pre-computed scores for all cds regions in the human genome.

Groups of KMT2D expression were generated by first normalizing all RNA-seq raw counts from the TCGA using DESeq2 (Love et al., 2014). Counts for KMT2D were then extracted, sorted, and grouped into quartiles. Clinical data from TCGA were right censored at the 2 year mark to generate Kaplan-Meier curves for 2-year disease-free survival in the KMT2D expression high vs. KMT2D expression low cohort.

### Cell Line RNA-seq

RNA extraction for HN30 knockdown experiments was performed using an RNeasy Mini Kit per manufacturer’s instructions (Qiagen). Isolation of mRNA was performed using NEBNext Poly(A) mRNA Magnetic Isolation Module and libraries were prepared using the NEBNext Ultra II Directional RNA Library Prep kit (New England BioLabs). Library quality was checked on an Agilent TapeStation 4150 and quantified by Qubit 2000 fluorometer (Invitrogen). Libraries were pooled in equimolar ratios and sequenced on Illumina NovaSeq6000 SP runs with paired-end 100-bp reads at the The Advanced Technology Genomics Core (ATGC) at MD Anderson Cancer Center.

RNA-seq raw reads were processed using the provided pipeline: https://github.com/sccallahan/QUACKERS_RNAseq-pipeline. In brief, raw reads were aligned to the hg19 genome using STAR v2.7.2b (Dobin et al., 2013) and quality checked using FastQC v0.11.8 (Bioinformatics) (https://www.bioinformatics.babraham.ac.uk/projects/fastqc/). Counts were generated using featureCounts from subread v1.6.3 (Liao et al., 2013). Downstream normalization and differential expression analysis were performed using DESeq2 (Love et al., 2014). Pathway enrichment analyses were performed using GSEA’s pre-ranked list option (Subramanian et al., 2005).

### Cell Line Whole Exome Sequencing (WES)

WES data was processed as previously described and obtained as a MAF file from the authors (Gleber-Netto et al., 2019; Subramanian et al., 2005). To cluster the cell lines based on mutation background, all mutation calls were binzarized to 1 or 0 to represent “mutated” or “not mutated,” respectively. The Jaccard distance matrix was then computed, and the resulting matrix was clustered using Ward’s minimum variance method.

### Chromatin Immunoprecipitation followed by Next Generation Sequencing (ChIP-Seq)

ChIP assays were performed as described previously (Terranova et al., 2018). Briefly, approximately 2 × 10^7^ cells were harvested by scraping. Samples were cross-linked with 1% (wt/ vol) formaldehyde for 10 min at 37 °C with shaking. After quenching with 150 mM glycine for 5 min at 37 °C with shaking, cells were washed twice with ice-cold PBS and frozen at -80 °C for further processing. Cross-linked pellets were thawed and lysed on ice for 30 min in ChIP harvest buffer (12 mM Tris-Cl, 1 × PBS, 6 mM EDTA, 0.5% SDS) with protease inhibitors (Sigma). Lysed cells were sonicated with a Bioruptor (Diagenode) to obtain chromatin fragments (∼200–500 bp) and centrifuged at 15,000 × g for 15 min to obtain a soluble chromatin fraction. In parallel with cellular lysis and sonication, antibodies (5 μg/3 × 10^6^ cells) were coupled with 30 μl of magnetic protein G beads in binding/blocking buffer (PBS + 0.1% Tween + 0.2% BSA) for 2h at 4 °C with rotation. Antibodies used for ChIP included anti-H3K4me3 (Abcam; ab8580), anti-H3K4me1 (Abcam; ab8895), and anti-H3K27ac (Abcam; ab4729). Soluble chromatin was diluted five times using ChIP dilution buffer (10 mM Tris-Cl, 140 mM NaCl, 0.1% DOC, 1% Triton X, 1 mM EDTA) with protease inhibitors and added to the antibody-coupled beads with rotation at 4 °C overnight. After washing, samples were treated with elution buffer (10 mM Tris-Cl, pH 8.0, 5 mM EDTA, 300 mM NaCl, 0.5% SDS), RNase A, and Proteinase K, and cross-links were reversed overnight at 37. Immune complexes were then washed five times with cold RIPA buffer (10mM Tris–HCl, pH 8.0, 1mM EDTA, pH 8.0, 140mM NaCl, 1% Triton X-100, 0.1% SDS, 0.1% DOC), twice with cold high-salt RIPA buffer (10mM Tris–HCl, pH 8.0, 1mM EDTA, pH 8.0, 500mM NaCl, 1% Triton X-100, 0.1% SDS, 0.1% DOC), and twice with cold LiCl buffer (10mM Tris– HCl, pH 8.0, 1mM EDTA, pH 8.0, 250mM LiCl, 0.5% NP-40, 0.5% DOC). ChIP DNA was purified using SPRI beads (Beckman Coulter) and quantified using the Qubit 2000 (Invitrogen) and TapeStation 4150 (Agilent). Libraries for Illumina sequencing were generated following the New England BioLabs (NEB) Next Ultra DNA Library Prep Kit protocol. Amplified ChIP DNA was purified using double-sided SPRI to retain fragments (∼200–500 bp) and quantified using the Qubit 2000 and TapeStation 4150 before multiplexing.

Raw fastq reads for all ChIP-seq experiments were processed using a snakemake based pipeline https://github.com/crazyhottommy/pyflow-ChIPseq. Briefly, raw reads were first processed using FastQC (https://www.bioinformatics.babraham.ac.uk/projects/fastqc/) and uniquely mapped reads were aligned to the hg19 reference genome using Bowtie version 1.1.2 (Langmead et al., 2009). Duplicate reads were removed using SAMBLASTER (Faust and Hall, 2014) before compression to bam files. To directly compare ChIP-seq samples, uniquely mapped reads for each mark were downsampled per condition to 15 million, sorted, and indexed using samtools version 1.5 (Li et al., 2009). To visualize ChIP-seq libraries on the IGV genome browser, we used deepTools version 2.4.0 (Ramirez et al., 2014) to generate bigWig files by scaling the bam files to reads per kilobase per million (RPKM).

Peak overlaps were performed using bedtools (Quinlan and Hall, 2010) and differential binding analyses were performed using DiffBind (Stark, 2020) (https://bioconductor.org/packages/release/bioc/html/DiffBind.html). Enhancer-promoter linkages were established using data from Cao et al (Cao et al., 2017), and pathway analyses were performed using the MSigDB webtool (http://www.gsea-msigdb.org/gsea/msigdb/index.jsp). Motif analyses were performed using the RSAT web tool NGS-ChIP-seq module (http://rsat.sb-roscoff.fr/).

### Tissue Microarrays

A TMA of human HNSCC core was commercially acquired from Biomax (TMA HN803). The TMA was stained with anti-KMT2D (Sigma-Aldrich HPA035977) and scored by a medical pathologist at the MD Anderson Center.

### Mouse studies

4-6 weeks old female mice on C57BL/6 background were used for *in vivo* studies. To induce mouse oral cavity tumor, doxycycline inducible shKMT2D knockdown and shNT MOC1 cell lines were implanted into the tongue with subsequent supplementation of doxycycline containing water (2 mg/mL). For IFN cytokine treatment experiment, mice were subjected to IFN administration via i.p. route in the amount of 1 ug per mouse every other day after tumor establishment. Mice were housed at the SPF conditions in the animal facility at M.D. Anderson cancer Center and all animal experiments were approved by the Institutional Animal Care and Use Committee.

### Immunoprofiling with flow cytometry analysis and ELISA

Tongue tumor, spleen and draining lymph nodes for flow cytometry analysis were harvested at the terminal point of the *in vivo* experiments. Single-cell suspension was utilized for subsequent FACS analysis. For intracellular and nuclear staining, cells were stimulated in the presence of phorbol 12-myristate-13-acetate, ionomycin, and brefeldin A for 4 hrs. Fixation and permeabilization buffers from eBioscience were used according to manufacturer’s instructions. Samples were acquired with LSR Fortessa X-20 flow cytometer (BD Bioscience) and analyzed with FlowJo software. For ELISA analysis serum isolated from terminal heart puncture and supernatant collected from activated overnight in the presence of 2.5 ug agonistic anti-CD3 and anti-CD28 antibodies and 1 ug/mL of LPS cells were used to detect IFN*γ*. Antibodies and reagents used for ELISA and Flow cytometry analysis are listed in the **Supplementary Table 2**.

## FIGURE LEGENDS

**FIGURE S1.**
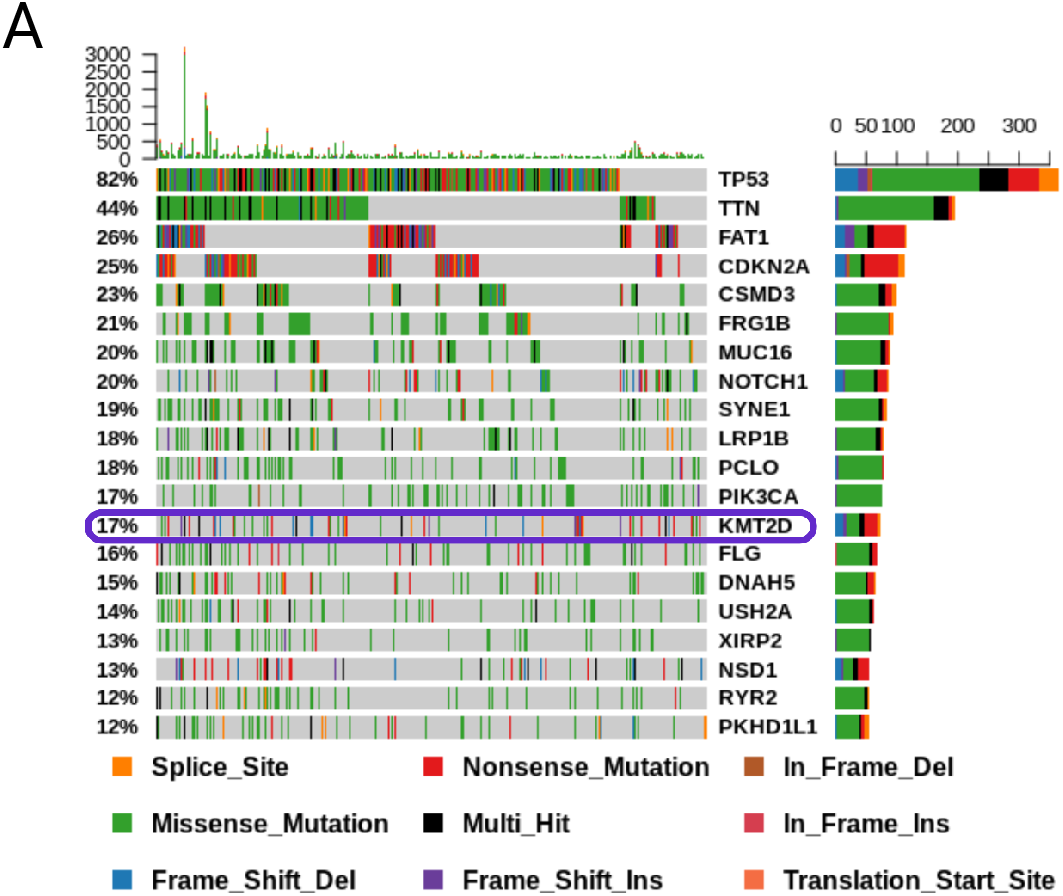

**Supplementary Table 1:** List of KMT2D WT and Mut Cell Lines Used in This Study.

| **SampleID** | **Tissue** | **Mut_type** |
| --- | --- | --- |
| MDA686TU | Mut | frameshift |
| MDA1586 | Mut | frameshift |
| JHU029 | Mut | stopgain |
| MDA686LN | Mut | frameshift |
| MDA886LN | Mut | stopgain |
| HN5 | WT | NA |
| PCI24 | WT | NA |
| UMSCC6 | WT | NA |
| UMSCC11A | WT | NA |
| UMSCC22B | WT | NA |

**Supplementary Table 2.**

| Reagent name | Catalog number | Vendor | Concentration/dilution used |
| --- | --- | --- | --- |
| CD45-BV711 | 563709 | BD Biosciences | 1:200 |
| CD8a-AF700 | 100729 | Biolegend | 1:200 |
| Live/dead marker -BV510 | 13-0870-T100 | Tonbo Biosciences | 1:500 |
| CD3-BV421 | 100213 | Biolegend | 1:200 |
| CD4-BV785 | 100453 | Biolegend | 1:200 |
| CD335-APC | 137607 | Biolegend | 1:200 |
| Foxp3-AF647 | 126407 | Biolegend | 1:100 |
| NK1.1-PerCPCy5.5 | 108727 | Biolegend | 1:200 |
| PD-1- PECF594 | 135227 | Biolegend | 1:200 |
| CTLA-4 Pecy7 | 106313 | Biolegend | 1:200 |
| CD62L-BV421 | 104423 | Biolegend | 1:200 |
| CD44-PE | 103007 | Biolegend | 1:200 |
| CD4-FITC | 100425 | Biolegend | 1:200 |
| Tim-3- PerCPCy5.5 | 134011 | Biolegend | 1:200 |
| IFNγ-BV421 | 505829 | Biolegend | 1:100 |
| Granzyme B-APC | 372203 | Biolegend | 1:100 |
| Ki-67-PECF594 | 652427 | Biolegend | 1:100 |
| Capture IFNγ | 505701 | Biolegend | 5 ug/mL |
| Detection IFNγ | 505803 | Biolegend | 1 ug/mL |
| HRP-Avidin D | A-2014 | Vector Laboratories | 1:2000 |
| TMB | 34028 | Thermo Scientific | according to manufacturer |
| Ionomycin calcium salt | I0634-1MG | Sigma-Aldrich | 1000x from 1.5 mM stock |
| Phorbol 12-myristate 13-acetate | P8139-1MG | Sigma-Aldrich | 0.1 ug/mL |
| Brefeldin A | 00-4506-51 | Life Technologies | 1000x |
| CD28 monoclonal antibody | 16-0281-81 | Life Technologies | 2.5 ug/sample |
| CD3 monoclonal antibody | 16-0031-81 | Life Technologies | 2.5 ug/sample |
| Permeabilization Buffer (10X) | 00-8333-56 | Life Technologies | according to manufacturer |
| Fixation/Permeabilization Diluent | 00-5223-56 | Life Technologies | according to manufacturer |
| Fixation/Permeabilization Concentrate | 00-5123-43 | Life Technologies | according to manufacturer |
| IC Fixation Buffer | 00-8222-49 | Life Technologies | according to manufacturer |
| Lipopolysaccharides | L2654-1MG | Sigma-Aldrich | 1 ug/mL |

## CITATIONS

Alam, H., Tang, M., Maitituoheti, M., Dhar, S.S., Kumar, M., Han, C.Y., Ambati, C.R., Amin, S.B., Gu, B., Chen, T.Y., et al. (2020). KMT2D Deficiency Impairs Super-Enhancers to Confer a Glycolytic Vulnerability in Lung Cancer. Cancer Cell 37, 599–617 e597.

Almeida, L.O., Abrahao, A.C., Rosselli-Murai, L.K., Giudice, F.S., Zagni, C., Leopoldino, A.M., Squarize, C.H., and Castilho, R.M. (2014). NFkappaB mediates cisplatin resistance through histone modifications in head and neck squamous cell carcinoma (HNSCC). FEBS Open Bio 4, 96–104.

Argiris, A., Karamouzis, M.V., Raben, D., and Ferris, R.L. (2008). Head and neck cancer. Lancet 371, 1695–1709.

Ayers, M., Lunceford, J., Nebozhyn, M., Murphy, E., Loboda, A., Kaufman, D.R., Albright, A., Cheng, J.D., Kang, S.P., Shankaran, V., et al. (2017). IFN-gamma-related mRNA profile predicts clinical response to PD-1 blockade. J Clin Invest 127, 2930–2940.

Baylin, S.B., and Jones, P.A. (2016). Epigenetic Determinants of Cancer. Cold Spring Harb Perspect Biol 8.

Bioinformatics, B. FastQC A Quality Control tool for High Throughput Sequence Data.

Bonner, J.A., Harari, P.M., Giralt, J., Azarnia, N., Shin, D.M., Cohen, R.B., Jones, C.U., Sur, R., Raben, D., Jassem, J., et al. (2006). Radiotherapy plus cetuximab for squamous-cell carcinoma of the head and neck. N Engl J Med 354, 567–578.

Briesemeister, D., Sommermeyer, D., Loddenkemper, C., Loew, R., Uckert, W., Blankenstein, T., and Kammertoens, T. (2011). Tumor rejection by local interferon gamma induction in established tumors is associated with blood vessel destruction and necrosis. Int J Cancer 128, 371–378.

Burtness, B., Harrington, K.J., Greil, R., Soulieres, D., Tahara, M., de Castro, G., Jr., Psyrri, A., Baste, N., Neupane, P., Bratland, A., et al. (2019). Pembrolizumab alone or with chemotherapy versus cetuximab with chemotherapy for recurrent or metastatic squamous cell carcinoma of the head and neck (KEYNOTE-048): a randomised, open-label, phase 3 study. Lancet 394, 1915–1928.

Cancer Genome Atlas, N. (2015). Comprehensive genomic characterization of head and neck squamous cell carcinomas. Nature 517, 576–582.

Cao, Q., Anyansi, C., Hu, X., Xu, L., Xiong, L., Tang, W., Mok, M.T.S., Cheng, C., Fan, X., Gerstein, M., et al. (2017). Reconstruction of enhancer-target networks in 935 samples of human primary cells, tissues and cell lines. Nature genetics 49, 1428–1436.

Castilho, R.M., Squarize, C.H., and Almeida, L.O. (2017). Epigenetic Modifications and Head and Neck Cancer: Implications for Tumor Progression and Resistance to Therapy. Int J Mol Sci 18.

Dhar, S.S., and Lee, M.G. (2021). Cancer-epigenetic function of the histone methyltransferase KMT2D and therapeutic opportunities for the treatment of KMT2D-deficient tumors. Oncotarget 12, 1296–1308.

Dobin, A., Davis, C.A., Schlesinger, F., Drenkow, J., Zaleski, C., Jha, S., Batut, P., Chaisson, M., and Gingeras, T.R. (2013). STAR: ultrafast universal RNA-seq aligner. Bioinformatics 29, 15–21.

Easwaran, H., Tsai, H.C., and Baylin, S.B. (2014). Cancer epigenetics: tumor heterogeneity, plasticity of stem-like states, and drug resistance. Mol Cell 54, 716–727.

ElKassar, N., and Gress, R.E. (2010). An overview of IL-7 biology and its use in immunotherapy. J Immunotoxicol 7, 1–7.

Faust, G.G., and Hall, I.M. (2014). SAMBLASTER: fast duplicate marking and structural variant read extraction. Bioinformatics 30, 2503–2505.

Ferris, R.L., Blumenschein, G., Jr., Fayette, J., Guigay, J., Colevas, A.D., Licitra, L., Harrington, K., Kasper, S., Vokes, E.E., Even, C., et al. (2016). Nivolumab for Recurrent Squamous-Cell Carcinoma of the Head and Neck. N Engl J Med 375, 1856–1867.

Forastiere, A., Koch, W., Trotti, A., and Sidransky, D. (2001). Head and neck cancer. N Engl J Med 345, 1890–1900.

Fridman, W.H., Pages, F., Sautes-Fridman, C., and Galon, J. (2012). The immune contexture in human tumours: impact on clinical outcome. Nat Rev Cancer 12, 298–306.

Froimchuk, E., Jang, Y., and Ge, K. (2017). Histone H3 lysine 4 methyltransferase KMT2D. Gene 627, 337–342.

Gleber-Netto, F.O., Rao, X., Guo, T., Xi, Y., Gao, M., Shen, L., Erikson, K., Kalu, N.N., Ren, S., Xu, G., et al. (2019). Variations in HPV function are associated with survival in squamous cell carcinoma. JCI Insight 4.

Gougis, P., Moreau Bachelard, C., Kamal, M., Gan, H.K., Borcoman, E., Torossian, N., Bieche, I., and Le Tourneau, C. (2019). Clinical Development of Molecular Targeted Therapy in Head and Neck Squamous Cell Carcinoma. JNCI Cancer Spectr 3, pkz055.

Gupta, P.B., Fillmore, C.M., Jiang, G., Shapira, S.D., Tao, K., Kuperwasser, C., and Lander, E.S. (2011). Stochastic state transitions give rise to phenotypic equilibrium in populations of cancer cells. Cell 146, 633–644.

India Project Team of the International Cancer Genome, C. (2013). Mutational landscape of gingivo-buccal oral squamous cell carcinoma reveals new recurrently-mutated genes and molecular subgroups. Nat Commun 4, 2873.

Janeway, C.A., Jr., and Medzhitov, R. (2002). Innate immune recognition. Annu Rev Immunol 20, 197–216.

Jefferies, C.A. (2019). Regulating IRFs in IFN Driven Disease. Front Immunol 10, 325.

Joyce, J.A., and Pollard, J.W. (2009). Microenvironmental regulation of metastasis. Nat Rev Cancer 9, 239–252.

Judd, N.P., Allen, C.T., Winkler, A.E., and Uppaluri, R. (2012a). Comparative analysis of tumor-infiltrating lymphocytes in a syngeneic mouse model of oral cancer. Otolaryngol Head Neck Surg 147, 493–500.

Judd, N.P., Winkler, A.E., Murillo-Sauca, O., Brotman, J.J., Law, J.H., Lewis, J.S., Jr., Dunn, G.P., Bui, J.D., Sunwoo, J.B., and Uppaluri, R. (2012b). ERK1/2 regulation of CD44 modulates oral cancer aggressiveness. Cancer Res 72, 365–374.

Kandoth, C., McLellan, M.D., Vandin, F., Ye, K., Niu, B., Lu, C., Xie, M., Zhang, Q., McMichael, J.F., Wyczalkowski, M.A., et al. (2013). Mutational landscape and significance across 12 major cancer types. Nature 502, 333–339.

Kantidakis, T., Saponaro, M., Mitter, R., Horswell, S., Kranz, A., Boeing, S., Aygun, O., Kelly, G.P., Matthews, N., Stewart, A., et al. (2016). Mutation of cancer driver MLL2 results in transcription stress and genome instability. Genes Dev 30, 408–420.

Kelley, D.Z., Flam, E.L., Izumchenko, E., Danilova, L.V., Wulf, H.A., Guo, T., Singman, D.A., Afsari, B., Skaist, A.M., Considine, M., et al. (2017). Integrated Analysis of Whole-Genome ChIP-Seq and RNA-Seq Data of Primary Head and Neck Tumor Samples Associates HPV Integration Sites with Open Chromatin Marks. Cancer Res 77, 6538–6550.

Klymenko, Y., and Nephew, K.P. (2018). Epigenetic Crosstalk between the Tumor Microenvironment and Ovarian Cancer Cells: A Therapeutic Road Less Traveled. Cancers (Basel) 10.

Koutsioumpa, M., Hatziapostolou, M., Polytarchou, C., Tolosa, E.J., Almada, L.L., Mahurkar-Joshi, S., Williams, J., Tirado-Rodriguez, A.B., Huerta-Yepez, S., Karavias, D., et al. (2019). Lysine methyltransferase 2D regulates pancreatic carcinogenesis through metabolic reprogramming. Gut 68, 1271–1286.

Langmead, B., Trapnell, C., Pop, M., and Salzberg, S.L. (2009). Ultrafast and memory-efficient alignment of short DNA sequences to the human genome. Genome biology 10, R25.

Lawson, A.R.J., Abascal, F., Coorens, T.H.H., Hooks, Y., O’Neill, L., Latimer, C., Raine, K., Sanders, M.A., Warren, A.Y., Mahbubani, K.T.A., et al. (2020). Extensive heterogeneity in somatic mutation and selection in the human bladder. Science 370, 75–82.

Leach, D.R., Krummel, M.F., and Allison, J.P. (1996). Enhancement of antitumor immunity by CTLA-4 blockade. Science 271, 1734–1736.

Lee, J.E., Wang, C., Xu, S., Cho, Y.W., Wang, L., Feng, X., Baldridge, A., Sartorelli, V., Zhuang, L., Peng, W., et al. (2013). H3K4 mono- and di-methyltransferase MLL4 is required for enhancer activation during cell differentiation. Elife 2, e01503.

Li, H., Handsaker, B., Wysoker, A., Fennell, T., Ruan, J., Homer, N., Marth, G., Abecasis, G., Durbin, R., and Genome Project Data Processing, S. (2009). The Sequence Alignment/Map format and SAMtools. Bioinformatics 25, 2078–2079.

Li, R., Du, Y., Chen, Z., Xu, D., Lin, T., Jin, S., Wang, G., Liu, Z., Lu, M., Chen, X., et al. (2020). Macroscopic somatic clonal expansion in morphologically normal human urothelium. Science 370, 82–89.

Liao, Y., Smyth, G.K., and Shi, W. (2013). The Subread aligner: fast, accurate and scalable read mapping by seed-and-vote. Nucleic Acids Res 41, e108.

Lin-Shiao, E., Lan, Y., Coradin, M., Anderson, A., Donahue, G., Simpson, C.L., Sen, P., Saffie, R., Busino, L., Garcia, B.A., et al. (2018). KMT2D regulates p63 target enhancers to coordinate epithelial homeostasis. Genes Dev 32, 181–193.

Liu, M., Guo, S., and Stiles, J.K. (2011). The emerging role of CXCL10 in cancer (Review). Oncol Lett 2, 583–589.

Local, A., Huang, H., Albuquerque, C.P., Singh, N., Lee, A.Y., Wang, W., Wang, C., Hsia, J.E., Shiau, A.K., Ge, K., et al. (2018). Identification of H3K4me1-associated proteins at mammalian enhancers. Nat Genet 50, 73–82.

Love, M.I., Huber, W., and Anders, S. (2014). Moderated estimation of fold change and dispersion for RNA-seq data with DESeq2. Genome Biol 15, 550.

Maitituoheti, M., Keung, E.Z., Tang, M., Yan, L., Alam, H., Han, G., Singh, A.K., Raman, A.T., Terranova, C., Sarkar, S., et al. (2020). Enhancer Reprogramming Confers Dependence on Glycolysis and IGF Signaling in KMT2D Mutant Melanoma. Cell Rep 33, 108293.

Marks, D.L., Olson, R.L., and Fernandez-Zapico, M.E. (2016). Epigenetic control of the tumor microenvironment. Epigenomics 8, 1671–1687.

Martinez-Jimenez, F., Muinos, F., Sentis, I., Deu-Pons, J., Reyes-Salazar, I., Arnedo-Pac, C., Mularoni, L., Pich, O., Bonet, J., Kranas, H., et al. (2020). A compendium of mutational cancer driver genes. Nat Rev Cancer 20, 555–572.

Mayakonda, A., Lin, D.C., Assenov, Y., Plass, C., and Koeffler, H.P. (2018). Maftools: efficient and comprehensive analysis of somatic variants in cancer. Genome Res 28, 1747–1756.

Medzhitov, R., and Janeway, C.A., Jr. (2002). Decoding the patterns of self and nonself by the innate immune system. Science 296, 298–300.

Mularoni, L., Sabarinathan, R., Deu-Pons, J., Gonzalez-Perez, A., and Lopez-Bigas, N. (2016). OncodriveFML: a general framework to identify coding and non-coding regions with cancer driver mutations. Genome Biol 17, 128.

Newman, A.M., Liu, C.L., Green, M.R., Gentles, A.J., Feng, W., Xu, Y., Hoang, C.D., Diehn, M., and Alizadeh, A.A. (2015). Robust enumeration of cell subsets from tissue expression profiles. Nat Methods 12, 453–457.

Nulton, T.J., Olex, A.L., Dozmorov, M., Morgan, I.M., and Windle, B. (2017). Analysis of The Cancer Genome Atlas sequencing data reveals novel properties of the human papillomavirus 16 genome in head and neck squamous cell carcinoma. Oncotarget 8, 17684–17699.

Onken, M.D., Winkler, A.E., Kanchi, K.L., Chalivendra, V., Law, J.H., Rickert, C.G., Kallogjeri, D., Judd, N.P., Dunn, G.P., Piccirillo, J.F., et al. (2014). A surprising cross-species conservation in the genomic landscape of mouse and human oral cancer identifies a transcriptional signature predicting metastatic disease. Clin Cancer Res 20, 2873–2884.

Pickering, C.R., Zhang, J., Yoo, S.Y., Bengtsson, L., Moorthy, S., Neskey, D.M., Zhao, M., Ortega Alves, M.V., Chang, K., Drummond, J., et al. (2013). Integrative genomic characterization of oral squamous cell carcinoma identifies frequent somatic drivers. Cancer Discov 3, 770–781.

Pitt, J.M., Marabelle, A., Eggermont, A., Soria, J.C., Kroemer, G., and Zitvogel, L. (2016). Targeting the tumor microenvironment: removing obstruction to anticancer immune responses and immunotherapy. Ann Oncol 27, 1482–1492.

Quinlan, A.R., and Hall, I.M. (2010). BEDTools: a flexible suite of utilities for comparing genomic features. Bioinformatics 26, 841–842.

Ramirez, F., Dundar, F., Diehl, S., Gruning, B.A., and Manke, T. (2014). deepTools: a flexible platform for exploring deep-sequencing data. Nucleic Acids Res 42, W187–191.

Rao, R.C., and Dou, Y. (2015). Hijacked in cancer: the KMT2 (MLL) family of methyltransferases. Nature reviews Cancer 15, 334–346.

Seif, F., Khoshmirsafa, M., Aazami, H., Mohsenzadegan, M., Sedighi, G., and Bahar, M. (2017). The role of JAK-STAT signaling pathway and its regulators in the fate of T helper cells. Cell Commun Signal 15, 23.

Stark, R., Brown, G. (2020). DiffBind: Differential Binding Analysis of ChIP-Seq Peak Data. Bioconductor version: Release (312).

Stransky, N., Egloff, A.M., Tward, A.D., Kostic, A.D., Cibulskis, K., Sivachenko, A., Kryukov, G.V., Lawrence, M.S., Sougnez, C., McKenna, A., et al. (2011). The mutational landscape of head and neck squamous cell carcinoma. Science 333, 1157–1160.

Stratton, M.R., Campbell, P.J., and Futreal, P.A. (2009). The cancer genome. Nature 458, 719–724.

Subramanian, A., Tamayo, P., Mootha, V.K., Mukherjee, S., Ebert, B.L., Gillette, M.A., Paulovich, A., Pomeroy, S.L., Golub, T.R., Lander, E.S., et al. (2005). Gene set enrichment analysis: a knowledge-based approach for interpreting genome-wide expression profiles. Proceedings of the National Academy of Sciences of the United States of America 102, 15545–15550.

Tamura, T., Yanai, H., Savitsky, D., and Taniguchi, T. (2008). The IRF family transcription factors in immunity and oncogenesis. Annu Rev Immunol 26, 535–584.

Tau, G.Z., Cowan, S.N., Weisburg, J., Braunstein, N.S., and Rothman, P.B. (2001). Regulation of IFN-gamma signaling is essential for the cytotoxic activity of CD8(+) T cells. J Immunol 167, 5574–5582.

Terranova, C., Tang, M., Orouji, E., Maitituoheti, M., Raman, A., Amin, S., Liu, Z., and Rai, K. (2018). An Integrated Platform for Genome-wide Mapping of Chromatin States Using High-throughput ChIP-sequencing in Tumor Tissues. J Vis Exp.

Toh, T.B., Lim, J.J., and Chow, E.K. (2017). Epigenetics in cancer stem cells. Mol Cancer 16, 29.

Veeramachaneni, R., Walker, T., Revil, T., Weck, A., Badescu, D., O’Sullivan, J., Higgins, C., Elliott, L., Liloglou, T., Risk, J.M., et al. (2019). Analysis of head and neck carcinoma progression reveals novel and relevant stage-specific changes associated with immortalisation and malignancy. Sci Rep 9, 11992.

Vermorken, J.B., Mesia, R., Rivera, F., Remenar, E., Kawecki, A., Rottey, S., Erfan, J., Zabolotnyy, D., Kienzer, H.R., Cupissol, D., et al. (2008). Platinum-based chemotherapy plus cetuximab in head and neck cancer. N Engl J Med 359, 1116–1127.

Wang, C., Lee, J.E., Lai, B., Macfarlan, T.S., Xu, S., Zhuang, L., Liu, C., Peng, W., and Ge, K. (2016). Enhancer priming by H3K4 methyltransferase MLL4 controls cell fate transition. Proc Natl Acad Sci U S A 113, 11871–11876.

Yanai, H., Negishi, H., and Taniguchi, T. (2012). The IRF family of transcription factors: Inception, impact and implications in oncogenesis. Oncoimmunology 1, 1376–1386.

Zhang, J., Dominguez-Sola, D., Hussein, S., Lee, J.E., Holmes, A.B., Bansal, M., Vlasevska, S., Mo, T., Tang, H., Basso, K., et al. (2015). Disruption of KMT2D perturbs germinal center B cell development and promotes lymphomagenesis. Nat Med 21, 1190–1198.

Zhao, M., Sano, D., Pickering, C.R., Jasser, S.A., Henderson, Y.C., Clayman, G.L., Sturgis, E.M., Ow, T.J., Lotan, R., Carey, T.E., et al. (2011). Assembly and initial characterization of a panel of 85 genomically validated cell lines from diverse head and neck tumor sites. Clin Cancer Res 17, 7248–7264.

Zhou, G., Liu, Z., and Myers, J.N. (2016). TP53 Mutations in Head and Neck Squamous Cell Carcinoma and Their Impact on Disease Progression and Treatment Response. J Cell Biochem 117, 2682–2692.

